# A multi-agent platform for assessment and improvement of bioinformatics software documentation

**DOI:** 10.64898/2026.02.09.704801

**Authors:** Anjun Ma, Shaohong Feng, Shaopeng Gu, Cankun Wang, Qin Ma

**Author notes:** Correspondence to: Anjun Ma.

## Abstract

Rapid advances in bioinformatics have transformed biomedical research in areas such as single-cell and spatial omics, digital pathology, and multi-modal data integration, yet software usability and reproducibility have not kept pace with the growing complexity and proliferation of computational tools. Inconsistent, incomplete, or inaccessible documentation remains a pervasive and underappreciated barrier, limiting tool adoption, hindering reproducibility across laboratories, and reducing the long-term impact of computational methods. Here, we introduce BioGuider, a multi-agent platform designed to systematically evaluate and improve documentation quality in bioinformatics software. Rather than treating documentation as ancillary text, BioGuider models it as a first-class, testable object. The platform implements a modular pipeline for documentation collection, assessment, reporting, and optional correction, with specialized agents that emulate real-world user interactions. BioGuider evaluates documentation against standardized, task-oriented criteria spanning installation, configuration, usage, and tutorials, and supports iterative, constraint-aware refinement while preserving code integrity and biological context. We benchmark BioGuider using a controlled error-injection framework that introduces realistic documentation failures across general, biology-specific, and configuration-related categories. Across multiple large language models, BioGuider demonstrates robust error detection and correction, with strong performance maintained under severe documentation degradation. Applying BioGuider to 47 widely used bioinformatics tools, we observe a positive association between documentation quality and citation frequency, highlighting documentation as a previously under-quantified driver of software adoption and scientific impact.

---

Over the past two decades, computational biology and bioinformatics have been fundamentally reshaped by rapid advances in software development^1^. High-throughput sequencing analytical tools have driven an unprecedented expansion. Unlike many general-purpose software engineering applications, bioinformatics software must routinely process heterogeneous data types, accommodate evolving experimental protocols, and encode specific biological assumptions. Analytical tasks are often multi-stage, parameter-rich, and sensitive to subtle preprocessing choices. Single-cell RNA sequencing (scRNA-seq) exemplifies these challenges. Some existing surveys suggest that more than 2,000 scRNA-seq tools have been developed^2^, many targeting quality control, normalization, batch correction, clustering, trajectory inference, cell– cell communication, spatial mapping, and cross-dataset integration. However, this rapid expansion has not been matched by equivalent gains in usability and accessibility. A systematic analysis by Mangul *et al*. estimated that 36,702 omics tools were published between 2005 and 2017, yet 28% were no longer accessible at the time of evaluation^3^. Among a randomly selected subset of 98 tools, 28% failed to install, and 57% failed to install when users followed only the provided installation tutorials. These observations underscore a persistent gap between methodological innovation and software usability and accessibility^4^. A dominant factor underlying this gap is documentation. Effective documentation should clearly define a tool’s purpose, scope, and assumptions, specify system requirements and dependencies, provide reliable installation instructions, and include executable examples that reflect realistic use cases. Achieving and maintaining this standard requires substantial time, expertise, and sustained effort, which are rarely incentivized in bioinformatics software development.

Large language models (LLMs) have demonstrated strong capabilities in code synthesis, refactoring, documentation drafting, and error explanation, while retrieval-augmented generation enables these models to ground their outputs in external knowledge sources such as code repositories and documentation files. Meanwhile, AI agents introduce a paradigm in which models can pursue defined objectives, interact with external environments, and iteratively refine their actions based on feedback. In industrial software engineering, this paradigm has motivated a growing body of systems designed to understand repository structure^5^, construct repository-level code graphs^6^, or automatically generate documentation by traversing codebases and dependency relationships^7^. However, these practices exhibit several limitations when considered in the context of bioinformatics software. First, many existing systems are developed as industrial-grade products or proprietary platforms and are not fully open-source or are not explicitly designed for bioinformatics workflows. Second, existing agent-based systems primarily focus on documentation generation or code understanding, but lack explicit, standardized criteria for evaluating documentation quality. As a result, they are not designed to assess or iteratively improve documentation for already published tools, nor to identify concrete failure modes that hinder installation, execution, or reuse. Third, the practical adoption of these systems often requires substantial setup effort, familiarity with agent orchestration frameworks, or integration into bespoke development pipelines, creating a high barrier for widespread use by tool developers and end users in the bioinformatics community.

To this end, we develop BioGuider, a multi-agent platform designed to evaluate and improve documentation in bioinformatics software. BioGuider is organized as a framework that reflects the end-to-end lifecycle of documentation usage and maintenance, comprising a collection module, an assessment module, a report module, and an optional document correction module (**Fig. 1a** and **Supplementary Methods**). Each module is implemented by specialized agents with clearly defined roles, enabling structured reasoning, execution, and feedback across heterogeneous software repositories. In the **collection module**, BioGuider first ingests documentation from GitHub repositories or local directories. A planning agent analyzes the repository structure and determines where relevant documentation artifacts are likely to reside. An execution agent then retrieves candidate files by traversing directories and loading documentation content, while a categorize agent classifies retrieved files into standardized documentation types, including README files, installation instructions, user guides or API references, and tutorials or vignettes. This module produces a curated and structured documentation corpus from repositories with diverse and often inconsistent layouts. The **assessment module** evaluates the collected documentation against predefined quality criteria. An optional testing agent executes installation procedures and tutorial workflows in controlled environments to assess practical reproducibility, capturing execution outcomes, error messages, and system-level failures. An assessment agent integrates these execution results with content-level analyses to generate structured evaluations that quantify documentation completeness, clarity, correctness, and consistency with the underlying codebase. In the **report module**, a reporting agent aggregates evaluation outputs into standardized quality control reports. These reports summarize strengths and weaknesses across documentation categories, link identified issues to specific files and sections, and provide concrete, evidence-based feedback that is consistent across repositories. Finally, a **correction module** enables iterative improvement of documentation (optional, depending on users’ requests). A dedicated correction agent translates evaluation feedback into targeted revisions of documentation text, structure, and examples. The module prompts are explicitly required for the correction of typographical, formatting, biological terminology, and configuration errors while enforcing strict constraints, including preservation of all code blocks, prohibition of content expansion, and prohibition of section removal.

**Fig 1.**
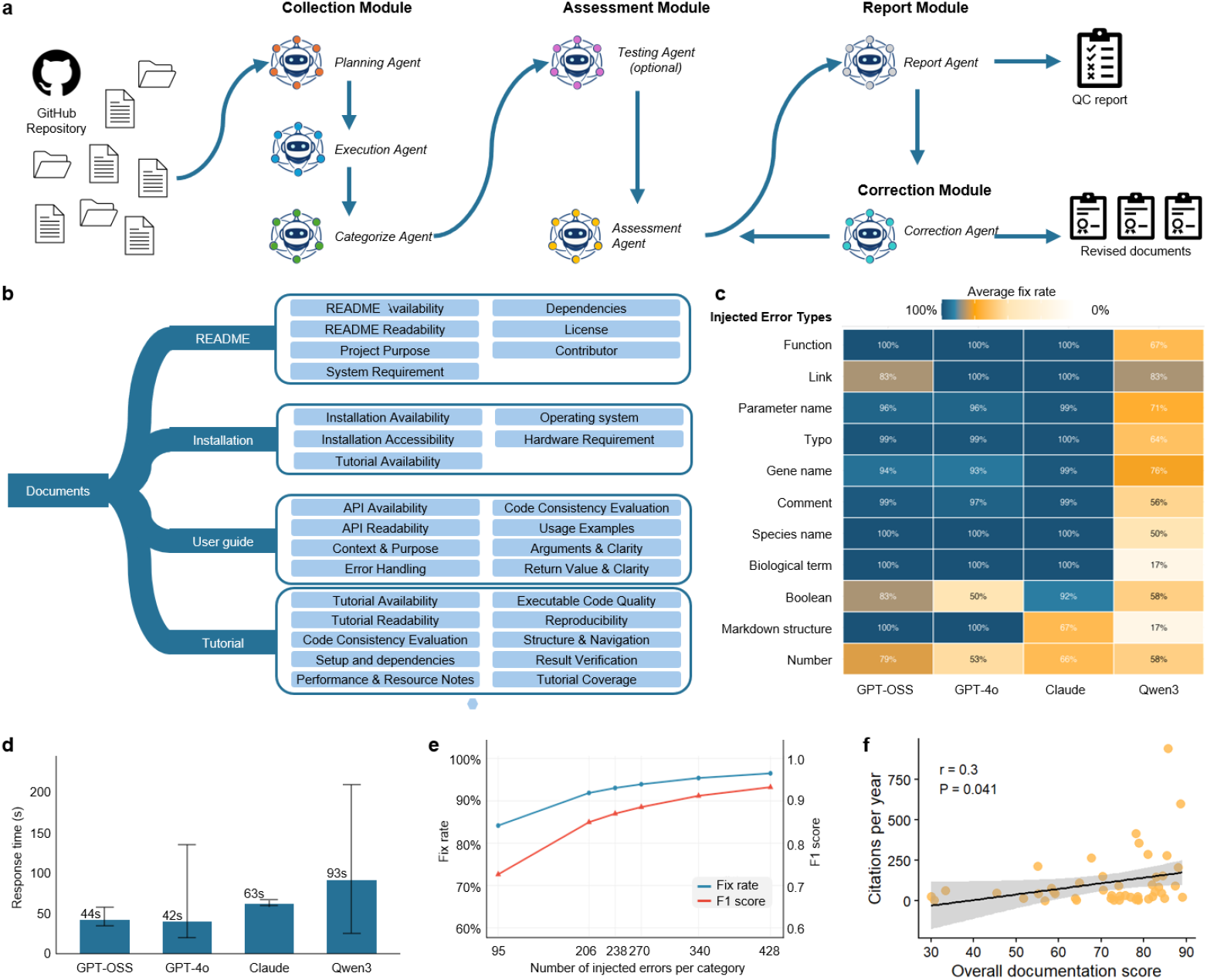
BioGuider framework, evaluation criteria, and benchmarking results. a. Overview of the BioGuider multi-agent framework for evaluating and improving bioinformatics software documentation. BioGuider is organized into four modules: a collection module that ingests documentation from GitHub repositories using planning, execution, and categorization agents; an assessment module that evaluates documentation quality and optionally tests reproducibility through controlled execution; a report module that summarizes evaluation results into standardized quality control reports; and an optional correction module that iteratively revises documentation based on identified deficiencies.b. Hierarchical documentation evaluation criteria implemented in BioGuider. Documentation is decomposed into four categories, each evaluated using category-specific metrics capturing availability, readability, clarity, reproducibility, consistency with source code, and usability-relevant features.c. Correction performance across different injected error types and large language models. Heatmap shows average fix rates for representative error categories spanning general documentation errors, biology-specific terminology errors, and command-line or configuration errors.d. Runtime comparison across evaluated language models, measured as average response time per document correction task.e. Stress testing of BioGuider under increasing documentation corruption. Fix rate and F1 score are shown as a function of the number of injected errors per document, demonstrating robustness under progressively degraded documentation conditions.f. Association between overall documentation quality and software impact. Scatter plot shows BioGuider overall documentation scores versus citation rates per year for 47 published bioinformatics software tools. The solid line indicates a linear fit with 95% confidence interval. Correlation statistics are shown in the panel. Citation statistics collected on Oct 30^th^, 2025.

A key design in BioGuider is a standardized evaluation framework that decomposes documentation quality into interpretable and task-oriented metrics (**Fig. 1b** and **Supplementary Table S1**). Rather than assigning a single undifferentiated score, BioGuider evaluates distinct documentation types independently, reflecting their different roles in the user experience. The design of this evaluation framework is informed by software publication guidelines and recommended best practices articulated across multiple bioinformatics-focused journals, which emphasize clarity, reproducibility, and usability but are often applied inconsistently in practice. README files are assessed based on availability, readability, clarity of project purpose, system requirements, dependency specification, licensing, and contributor information. Installation documentation is evaluated with emphasis on accessibility, operating system compatibility, hardware requirements, dependency complexity, and whether installation can be completed by following the documented instructions alone. User guides and API documentation are assessed for API availability, readability, contextual explanation, argument clarity, error handling, return value interpretation, usage examples, and consistency with the underlying code. Tutorials are evaluated using the most stringent criteria, including executability, reproducibility, setup completeness, performance considerations, structural navigation, result verification, and coverage of core functionalities. These metrics are designed to reflect real-world failure modes encountered by users.

To quantitatively evaluate BioGuider’s ability to detect and correct documentation errors, we established a controlled benchmarking framework based on systematic error injection into otherwise clean software documentation (**Supplementary Methods**). Starting from curated baseline documents, synthetic errors were introduced across multiple categories while preserving overall document structure, semantic coherence, and code block integrity. Error injection was performed using a constraint-based language model prompt that enforced minimum error counts per category, maintained high token-level overlap with the original text, and strictly preserved all code fence boundaries. To ensure that injected errors reflected realistic failure modes encountered in bioinformatics documentation, project-specific terminology was automatically extracted from source code files. Only corrupted documents were subsequently processed by BioGuider without access to the baseline document or the injected error manifest. We benchmarked multiple LLMs, including GPT-OSS, GPT-4o, Claude Sonnet, and Qwen3, with each model receiving identical corrupted inputs and correction prompts. GPT-OSS consistently achieved the highest correction performance across most error categories (**Fig. 1c**). In terms of runtime, GPT-OSS and GPT-4o exhibited comparable execution times, while GPT-OSS provided more stable correction behavior across heterogeneous error profiles (**Fig. 1d**). Based on this combined assessment of accuracy and efficiency, GPT-OSS was selected as the default language model for BioGuider implementation. Finally, we conducted stress testing by progressively increasing the number and density of injected errors per document to evaluate robustness under increasingly degraded documentation conditions. BioGuider maintained high correction performance even at elevated error loads, demonstrating its capacity to operate effectively on heavily corrupted and heterogeneous documentation (**Fig. 1e**). These results indicate that BioGuider can support reliable assessment and correction not only for well-maintained repositories but also for highly fragmented or legacy bioinformatics software documentation.

We next applied BioGuider to systematically evaluate the documentation quality of 47 published bioinformatics tools and examined whether documentation quality is associated with downstream usage and scientific impact (**Supplementary Table S2**). For each tool, BioGuider generated an overall documentation score by aggregating category-specific evaluations, which was then compared with citation rates normalized as citations per year. Across the evaluated tools, we observed a positive association between documentation quality and citation rate, with tools receiving higher BioGuider scores tending to accumulate citations more rapidly (**Fig. 1f**). Although substantial variability was present, reflecting the multifactorial nature of software adoption, the overall trend indicates that documentation quality is a meaningful contributor to software impact. These results suggest that well-documented tools are more likely to be discoverable, installable, and reusable by the broader community, thereby facilitating adoption beyond the original development context. Importantly, this analysis does not imply a direct causal relationship between documentation quality and citation counts, as software impact is influenced by many factors, including biological relevance, methodological novelty, and timing of release. Rather, the observed correlation highlights documentation as a previously under-quantified but practically important dimension of software sustainability and reuse.

To facilitate broad adoption and practical use, BioGuider is deployed through a web-based application programming interface that abstracts the underlying multi-agent workflow into a streamlined user experience (**Supplementary Note**). Developers can initiate a BioGuider evaluation by simply providing the GitHub repository URL of a software project, without requiring local installation or configuration. Upon completion of the evaluation, the platform generates a standardized quality control report that can be directly downloaded, summarizing documentation strengths, deficiencies, and category-specific scores. For users seeking iterative improvement, BioGuider further supports documentation correction workflows driven by the evaluation results. Corrected documentation files can be generated within the same interface, followed by a subsequent round of automated quality assessment to quantify improvements. Both the revised documents and their updated quality control reports are presented through the web interface, enabling transparent comparison between original and corrected documentation.

In summary, the usability, accessibility, and reproducibility of bioinformatics software increasingly hinge on documentation quality. Usually, bioinformatics tools must encode domain-specific biological assumptions, complex data preprocessing steps, evolving experimental conventions, and tightly coupled computational and biological interpretations. In this context, incomplete or inaccurate documentation directly impedes correct usage, reproducibility across datasets and laboratories, and long-term software sustainability. BioGuider demonstrates that documentation evaluation for bioinformatics software can be formalized, automated, and scaled through a multi-agent framework grounded in realistic analytical workflows. By systematically identifying documentation failure modes in bioinformatics pipelines and enabling iterative, constraint-aware corrections, BioGuider improves software accessibility, promotes correct usage, and enhances reproducibility. By elevating documentation to a first-class object of evaluation and improvement, BioGuider offers a practical path toward narrowing the gap between methodological innovation and reliable, reproducible bioinformatics research. Future extensions will expand coverage to additional bioinformatics documentation modalities, support longitudinal tracking of documentation evolution across software versions, and facilitate community-driven standards tailored to biological data analysis.

## Code Availability

The online BioGuider platform is accessible via https://bmblx.bmi.osumc.edu/bioguider.

## Acknowledgements

A.M is supported by National Institutes of Health R21DK140693, P01AI177687, and P01CA278732. Q.M.is supported in part by the NIH R01GM152585, U54AG075931, and R01DK138504. This work was also supported by the Pelotonia Institute of Immuno-Oncology. The content is solely the responsibility of the authors and does not necessarily represent the official views of the Pelotonia Institute of Immuno-Oncology or the NIH.

## Author Contributions

A.M. conceptualized the work. S.F. and C.W. designed the framework and developed the software. S.G and S.F. designed the evaluation criteria and performed an example tool test. C.W. performed benchmark and stress tests. A.M. and Q.M. drafted the manuscript, and all authors edited the manuscript.

## Ethics Declarations

*The authors declare no competing interests*

