## Supplementary Notes for "A multi-agent platform for assessment and improvement of bioinformatics software documentation"

**Table of Contents**

- Supplementary Methods
- Supplementary Note: BioGuider User Instruction and Tutorial
- Supplementary Table S1: Evaluation Criteria Description
- Supplementary Table S2: Tested 47 Bioinformatics Tools and BioGuider Scores

**Supplementary Methods**

1. **Multi-agent documentation evaluation framework**

BioGuider implements a modular, multi-agent system for systematic evaluation and improvement of documentation in open-source bioinformatics software repositories. The framework decomposes documentation evaluation into four coordinated modules: (i) Collection, (ii) Assessment, (iii) Report, and (iv) Correction. Each module is executed by specialized large language model agents equipped with domain-specific tools and operating under explicit constraints.

1.1. **Collect Module**

The Collect Module identifies, retrieves, and structures repository files relevant to documentation evaluation. Its primary objective is to assemble a curated corpus of documentation artifacts, including README files, installation instructions, user guides and API references, and tutorials or vignettes, from heterogeneous repository layouts. The Collect Module consists of three specialized agents operating in a plan-execute-verify loop.

**Design Agent** is responsible for high-level planning of the collection process. Given an evaluation objective, the agent analyzes repository metadata and generates a structured action plan that specifies which directories to explore, which file types to prioritize, and which documentation artifacts to extract. Typical planned actions include quantifying repository composition, identifying candidate documentation directories, summarizing documentation files, and locating executable tutorial or vignette content embedded in notebooks or scripts.

**Execute Agent** carries out the action plan produced by the Design Agent. It invokes the specified tools in sequence and records intermediate outputs, including extracted file contents, summaries, and execution results. To support flexible and repository-agnostic exploration, BioGuider provides the following tools to LLM agents: a Directory Reader that traverses the repository structure to enumerate files and subdirectories; a File Reader that loads the full content of text-based files (e.g., Markdown, R Markdown, Python, R source files); a Relevance Classifier that determines whether a file is relevant to a target documentation category; a Content Summarizer that produces concise semantic summaries of documentation files; a Content Extractor that extracts specific sections from larger documents; and a Python AST REPL Tool that executes Python code in a controlled environment to perform repository-level analyses.

**Categorize Agent** evaluates whether the collection objective has been satisfied. It inspects the outputs produced by the Execute agent to determine whether all required documentation categories have been adequately covered. If documentation artifacts are missing, incomplete, or ambiguous, the Observe agent generates corrective feedback that triggers additional planning cycles by the Design agent, enabling iterative refinement of the collection process.

The output of the Collection module is a structured, category-labeled documentation corpus that serves as input to the Assessment module.

1.2. **Assessment Module**

The Assessment module evaluates the collected documentation artifacts against predefined quality criteria, with an emphasis on completeness, clarity, reproducibility, and technical correctness. The module is implemented using two coordinated agents with complementary roles: a Testing agent, which evaluates practical reproducibility through execution, and an Assessment agent, which performs structured, category-specific documentation evaluation.

**Testing agent** assesses documentation-supported reproducibility by automatically building and executing a minimal “get-started” workflow in a containerized environment. This process evaluates whether documented instructions can be followed to successfully install and run the software.

The Testing agent is implemented as a plan–execute–observe loop:

- **Plan step:** Constructs a tool-level execution plan based on repository structure, available documentation files, and intermediate error messages. Planned actions may include extracting runnable code from notebooks, creating or correcting a minimal demonstration script, and generating a Docker file as the final execution artifact.
- **Execute step:** Executes the plan strictly using only the specified tools, including *extract_python_file_from_notebook*, *write_file*, and *generate_Dockerfile*. The agent records tool outputs and the path of the generated Docker file.
- **Observe step:** Runs docker build and docker run on the generated Docker file, captures build-time and runtime errors, and returns structured observations. When execution fails, the observed errors are used to trigger replanning. When execution succeeds, the final Docker file content is returned as the reproducibility artifact.

Several key constraints are enforced in this pipeline. Plans are regenerated only in response to observed build or runtime failures. Docker file generation is required to be the final planned action, ensuring that the final artifact reflects all applied fixes. Execution does not improvise and strictly follows the planned tool sequence.

**Assessment agent** evaluates documentation content independently of execution by applying standardized, category-specific evaluation criteria. Documentation artifacts are evaluated separately for each documentation type, reflecting their distinct roles in the user experience.

For each documentation category, the Assessment agent applies a consistent evaluation pipeline:

1. **File normalization and sanitation**
   Binary files are skipped, oversized files are excluded, HTML content is converted to plain text, and notebooks are reduced to markdown and executable code blocks.
2. **Readability analysis**
   Standard readability metrics are computed, including Flesch Reading Ease, Flesch–Kincaid Grade Level, Gunning Fog Index, and SMOG Index. These metrics are provided as auxiliary signals to the evaluator.
3. **Structured evaluation**
   An LLM generates a schema-constrained evaluation that scores category-specific criteria and produces targeted improvement suggestions.
4. **Free-form evaluation**
   For README and Installation documentation, a second LLM pass expands structured outputs into detailed, human-readable feedback with quoted text snippets and explanatory comments.
5. **Score aggregation**
   Sub-scores are combined via weighted aggregation to produce category-level scores and an overall documentation score.

Each documentation category is evaluated using tailored criteria aligned with real-world usage and common software publication guidelines:

**README:** Availability, readability, clarity of project purpose, hardware and software requirements, dependency specification, license information, and contributor or maintainer details.

**Prompt:** You are an expert in evaluating the quality of README files in software repositories.

```
Your task is to analyze the provided README file and generate a structured quality assessment based on the following criteria.
If a LICENSE file is present in the repository, its content will also be provided to support your evaluation of license-related criteria.
You must provide the evaluation score in your response.
---
### Evaluation Criteria
1. Available: Is the README accessible and present? Output: Yes or No
2. Readability: Evaluate based on readability metrics AND identify specific errors/issues in the text.
   - You must identify and list ALL errors and anomalies (typos, malformed links, markdown errors, image syntax errors, domain term errors, inconsistencies, formatting issues, and other anomalies).
   - For each error, provide the exact text snippet, error type, suggested correction, and explanation.
3. Project Purpose: Is the project's goal or function clearly stated? Output: Yes or No
4. Hardware and Software Requirements: Are hardware/software specs and compatibility details included?
5. Dependencies: Are all necessary software libraries and dependencies clearly listed?
6. License Information: Is license type clearly indicated?
7. Author/Contributor Info: Are contributor or maintainer details provided?
8. Overall Score: Give an overall quality rating of the README.
---
### Readability Metrics
Flesch Reading Ease: {flesch_reading_ease}
Flesch-Kincaid Grade Level: {flesch_kincaid_grade}
Gunning Fog Index: {gunning_fog_index}
SMOG Index: {smog_index}
---
### README Path
{readme_path}
### README Content
{readme_content}
### LICENSE Path
{license_path}
### LICENSE Summarized Content
{license_summarized_content}
```

- **Installation:** Presence and accessibility of installation instructions, dependency specification and complexity, operating system compatibility, hardware requirements, and whether installation can be completed by following the documented steps alone.

**Prompt:** You are an expert in evaluating the quality of installation information in software repositories.

```
Your task is to analyze the provided files related to installation and generate a structured quality assessment based on the following criteria.
---
### Evaluation Criteria
1. Installation Available: Is the installation section in document (like README.md or INSTALLATION)?
2. Installation Tutorial: Is the step-by-step installation tutorial provided?
3. Number of required Dependencies Installation: The number of dependencies required to install.
4. Compatible Operating System: Is the compatible operating system described?
5. Hardware Requirements: Are hardware requirements described?
6. Overall Score: Give an overall quality rating of the installation information.
---
### Installation Files Provided
{installation_files_content}
```

- **User Guide / API documentation:** Readability, contextual explanation of functionality, clarity of arguments and return values, error handling guidance, coverage of usage examples, and consistency with the underlying code.

**Prompt:** You are an expert in evaluating the quality of user guide in software repositories.

```
Your task is to analyze the provided files related to user guide and generate a structured quality assessment based on the following criteria.
---
1. Readability AND Error Detection:
   - Use Flesch Reading Ease, Flesch-Kincaid Grade, Gunning Fog, SMOG.
   - You must scan for and identify ALL error instances (typos, malformed links, markdown/RMarkdown errors, bio term errors, function name errors, inline code formatting errors, and other anomalies).
   - List each occurrence separately; do not group similar errors.
2. Arguments and Clarity: describe arguments and their usage with concrete improvement suggestions.
3. Return Value and Clarity: describe return values and meaning with improvement suggestions.
4. Context and Purpose: describe context and purpose with improvement suggestions.
5. Error Handling: describe error handling with improvement suggestions.
6. Usage Examples: describe usage examples with improvement suggestions.
7. Overall Score: output 0-100.
---
### User Guide Content
{userguide_content}
```

- **Tutorial / Vignette:** Readability, setup and dependency completeness, executability, reproducibility, structure and navigation, result verification, performance or resource considerations, and coverage of core functionalities.

**Prompt:** You are an expert in evaluating the quality of tutorials in software repositories.

```
Your task is to analyze the provided tutorial file and generate a structured quality assessment based on the following criteria.
---
1. Readability AND Error Detection:
   - Use Flesch Reading Ease, Flesch-Kincaid Grade, Gunning Fog, SMOG.
   - You must scan for and identify ALL error instances (typos, malformed links, markdown/RMarkdown errors, bio term errors, function name errors, inline code formatting errors, and other anomalies).
   - List each occurrence separately; do not group similar errors.
2. Coverage: whether it covers major steps, dependencies, prerequisites, setup, and example usage.
3. Reproducibility: whether it provides a clear description of reproducibility.
4. Structure and Navigation: logical sections, TOC/anchors, time estimates.
5. Executable Code Quality: executable and idiomatic code, no hard-coded paths.
6. Result Verification: expected outputs and acceptance criteria.
7. Performance and Resource Notes: CPU/GPU usage, memory, runtime estimates.
---
### Tutorial File Content
{tutorial_file_content}
```

For User Guides/APIs and Tutorials/Vignettes, the Assessment agent additionally evaluates consistency between documentation and source code through a two-step process: **(1)** BioGuider scans repository source code to build a structured index capturing function and class names, argument signatures, return values, and inline documentation. **(2)** Documented code usage is compared against the indexed code structure to verify that referenced functions or classes exist, argument names and ordering are correct, and documented behavior aligns with source-level definitions.

To enhance transparency and reproducibility, representative prompt templates used by the Assessment agent are provided, including structured evaluation prompts for README, Installation, User Guide/API, and Tutorial documentation. These prompts specify explicit evaluation criteria, required error detection behavior, scoring formats, and constraints on content modification.

1.3. **Report Module**

The Report module is responsible for converting structured outputs generated by the Assessment module into standardized, human-readable quality control reports. This module is implemented by a dedicated Report agent, whose primary objective is to ensure that documentation evaluation results are presented in a consistent, interpretable, and auditable format across repositories.

**Report agent** consumes multiple sources of structured information:

- **Assessment outputs**, provided as JSON objects containing per-file scores, identified issues, and targeted improvement suggestions generated by the Assessment agent
- **Optional testing outputs**, including Docker build and runtime results, execution logs, and error messages produced by the Testing agent
- **Optional metadata**, such as repository URL, commit SHA, language model identifier, and evaluation timestamp

These inputs allow the Report agent to integrate both content-level and execution-level signals into a unified report.

The Report agent generates documentation quality control reports through a multi-step processing pipeline:

- Assessment JSON outputs are parsed into a common internal schema. Required fields are validated, and missing or malformed entries are flagged to ensure report completeness and structural consistency.
- Evaluation results are grouped by documentation category (README, Installation, User Guide/API, Tutorial/Vignette) and further organized at the file level. Category-level summaries and overall scores are computed from aggregated results.
- The agent produces concise narrative summaries that highlight major strengths and deficiencies. Where applicable, summaries are supported by concrete evidence snippets extracted from the original documentation, enabling direct inspection of identified issues.

To ensure transparency and reproducibility, the Report module enforces explicit traceability between reported findings and source artifacts. Each identified issue is linked to its originating file and, when applicable, to specific sections within that file. Generated reports include model and version identifiers, evaluation configuration parameters, and timestamps. All intermediate JSON artifacts and final rendered reports are stored together to support auditing, comparison across evaluations, and longitudinal tracking.

1.4. **Correction module**

The Correction module provides an optional mechanism for iterative improvement of documentation based on outputs generated by the Assessment and Report modules. This module is implemented by a dedicated Correction agent and is activated only when users explicitly choose to optimize documentation using BioGuider. The primary objective of the Correction module is to address identified documentation issues while preserving code integrity, document structure, and domain-specific content.

The **Correction agent** operates exclusively on information produced by upstream modules, including structured assessment results and standardized quality control reports. For each documentation file selected for correction, the agent receives file-scoped issues, category-specific scores, and targeted improvement suggestions. The agent does not introduce new content beyond the scope of identified deficiencies and does not modify source code files. Documentation correction is performed through a constrained, multi-step workflow:

- **Issue extraction:** The agent parses assessment and report outputs to extract actionable issues and improvement suggestions associated with specific files and documentation categories.
- **Edit planning:** Based on the extracted issues, the agent generates a structured edit plan that maps each issue to a concrete modification operation. Planned edits are designed to be minimal, targeted, and non-overlapping, avoiding unnecessary rewriting or stylistic drift.
- **Document revision:** The agent applies the planned edits to generate revised documentation files. Throughout this process, strict constraints are enforced to preserve all code blocks, command-line examples, configuration snippets, and YAML frontmatter exactly as written unless explicitly identified as erroneous.

Upon completion of documentation revision, the updated files are automatically passed through the Assessment and Report modules to generate a second round of evaluation and quality control reports. This closed-loop design enables direct comparison between pre-correction and post-correction documentation scores and issue profiles, allowing users to quantitatively assess the impact of the applied changes.

All revisions produced by the Correction module are recorded with file-level change manifests to support traceability. Original and revised documents, along with their corresponding assessment and report artifacts, are stored together to enable auditing and longitudinal comparison. By design, the Correction module prioritizes conservative, evidence-driven edits and avoids speculative modifications.

**2. Error injection benchmark framework**

To systematically evaluate the capacity of large language models to detect and correct documentation errors, we developed a controlled error injection framework. The framework operates on clean baseline documentation files and introduces synthetic errors across multiple categories while preserving the overall document structure and readability. Error injection was performed using GPT-4 (Azure OpenAI) with a constraint-based prompt that enforced minimum error counts per category, maintained at least 85% token overlap with the original document, and preserved all code block delimiters exactly. When the language model output failed validation checks, the system automatically applied a deterministic fallback injection method to ensure reproducible error patterns.

The injection process extracted project-specific terminology from source code files using regular expression patterns to identify function names, class definitions, and frequently occurring technical terms. These extracted terms were prioritized as targets for function name misspellings and parameter errors, ensuring that injected errors reflected realistic mistakes specific to the software package under evaluation. Errors were organized into three primary domains encompassing 30 distinct categories. General documentation errors included typographical mistakes (spelling, grammar, and punctuation), malformed hyperlinks (missing scheme components, stray spaces, or incorrect domains), duplicated text fragments, broken markdown structure (header levels, list indentation, and code fence formatting), image syntax errors, inline code formatting issues (missing backticks), emphasis marker problems, table alignment errors, and incorrect code language tags in fenced blocks.

Biology-specific errors targeted domain knowledge accuracy and included gene symbol case alterations (e.g., BRCA1 to brca1), species terminology confusion (human versus mouse, GRCh38 versus mm10), reference genome mismatches, sequencing modality conflation (RNA-seq versus ATAC-seq versus proteomics), normalization terminology misuse (CPM, TPM, CLR, log1p), UMI versus read count confusion, batch effect terminology errors, quality control threshold mistakes (e.g., mitochondrial percentage 0.5 instead of 5), file format confusion (FASTQ, BAM, MTX, H5AD, RDS), strandedness claims, coordinate system errors (0-based versus 1-based indexing), unit and scale mistakes, sample type conflation (primary tissue versus cell line), and contamination terminology misuse (ambient RNA versus doublets). Command-line and configuration errors comprised parameter name misspellings, incorrect default value statements, and subtle path typographical errors.

Corrupted documents were processed through each model using a standardized correction prompt. The prompt instructed the model to act as an expert document proofreader for bioinformatics documentation, with explicit instructions to fix all typographical errors, truncated words, misspellings, broken URLs, gene and protein name capitalization, species names, markdown formatting issues, YAML frontmatter errors, and boolean or numeric value mistakes. Critical constraints required preservation of all code blocks exactly as written, prohibition of new content addition, and prohibition of section removal. Models were provided only the corrupted document text without access to the error manifest or baseline document.

Correction performance was quantified using precision, recall, and F1 score calculated from error-level classification. Each injected error was classified as a true positive if the error was successfully corrected (the mutated text was removed or replaced with valid content), or as a false negative if the error remained in the output document. False positives were detected through semantic analysis of document changes using a separate language model evaluation pass that identified harmful unintended modifications beyond the injected error corrections. Precision was calculated as the number of true positives divided by the sum of true positives and false positives. Recall (also reported as fix rate) was calculated as the number of true positives divided by the sum of true positives and false negatives. The F1 score was computed as the harmonic mean of precision and recall.

Category-specific evaluation logic was implemented to account for the diverse nature of error types. Typographical errors were marked as fixed if the original text was restored or the mutated text was removed with content rewriting. Link errors required restoration of well-formed markdown link syntax. Duplicate errors required reduction in the count of the duplicated snippet. Markdown structure errors required correction of malformed headers or spacing issues. Biology-specific term errors and function name errors required complete removal of the mutated snippet from the output text.

For the model comparison benchmark, four models were evaluated: GPT-OSS, GPT-4o, Claude Sonnet, Qwen3, and. Each model received identical corrupted input documents and correction prompts to ensure fair comparison.

3. **Stress test design**

The benchmark employed a stress testing methodology with progressively increasing error density to characterize model performance degradation under challenging conditions. Five error injection levels were tested: 10, 30, 50, and 100 errors per category, corresponding to total injected error counts ranging from 95 to 428 errors per document depending on the applicable categories for each file. Target documents were selected from the Seurat R package repository (satijalab/seurat), a widely used single-cell RNA sequencing analysis toolkit. The test corpus included README files, RMarkdown tutorial vignettes, and installation documentation. For each error level, the benchmark pipeline performed the following sequence: copying the baseline repository, injecting errors into selected files with manifest generation, executing model-based correction, evaluating corrections against the error manifest, and aggregating metrics across files and categories.

Execution utilized parallel processing with ThreadPoolExecutor for input/output-bound language model API calls during both injection and correction phases. Results were exported in JSON format with complete error-level details and CSV format for aggregate statistics suitable for visualization and statistical analysis. Each benchmark run recorded total duration, per-file metrics, per-category breakdown, and detailed error-by-error classification status.

**Supplementary Note: BioGuider User Instruction and Example**

To evaluate a GitHub repository, users provide the repository URL in the input field and initiate the analysis by clicking the **“Generate/Evaluate Documentation”** button. BioGuider then automatically executes the documentation evaluation pipeline. Upon completion, the generated evaluation report is rendered in the results panel located directly below the input area, allowing users to review the assessment outcomes interactively.

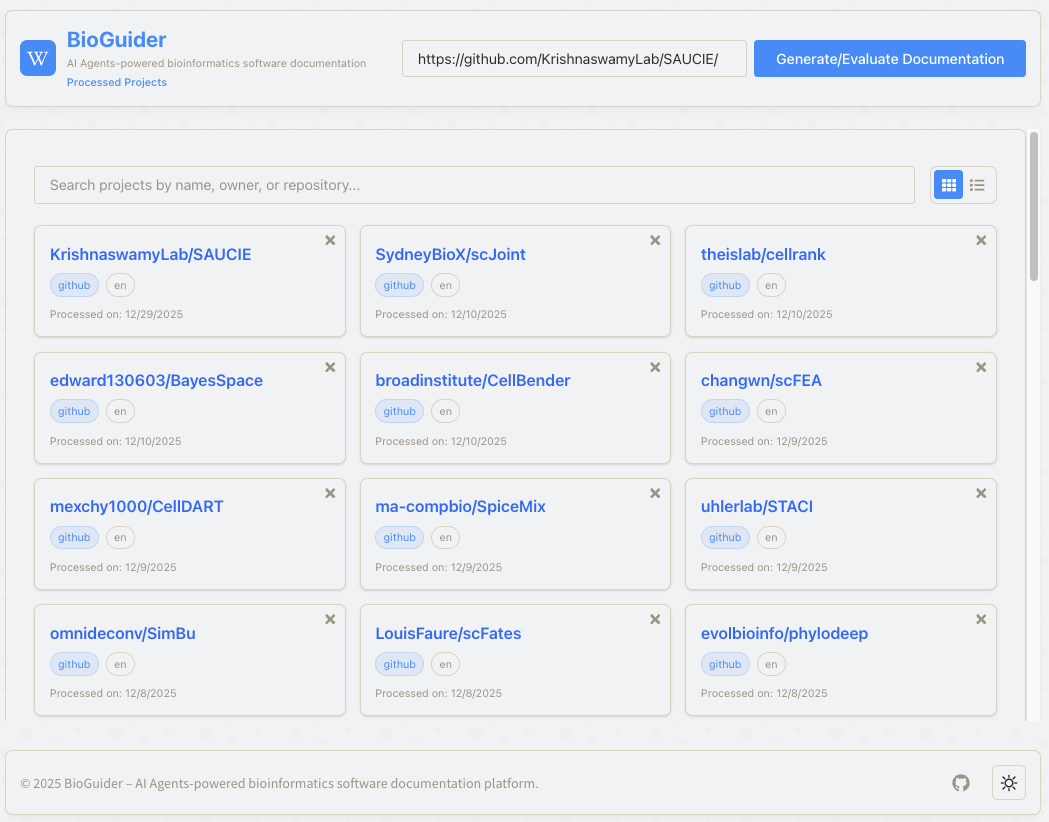

2 Click the button

1 Provide your GitHub repository link

Your evaluation history

Here, we used SAUCIE (<https://github.com/KrishnaswamyLab/SAUCIE/>), which was published in *Nature Methods* in 2019 (<https://doi.org/10.1038/s41592-019-0576-7>). SAUCIE is a deep neural network that combines parallelization and scalability to perform many single-cell data analysis tasks. In this document, we evaluated and refined the documents of SAUCIE to demonstrate the BioGuider’s functions.

After clicking the **“Generate/Evaluate Documentation”** button, a Configure dialog is displayed. This dialog allows users to specify evaluation parameters, including the documentation language and the large language model (LLM) provider and model to be used for analysis. BioGuider supports multiple LLM backends and model configurations, enabling users to tailor the evaluation process according to availability, cost, or performance considerations. Once the configuration is finalized, the evaluation is initiated by clicking **“Generate Report”**.

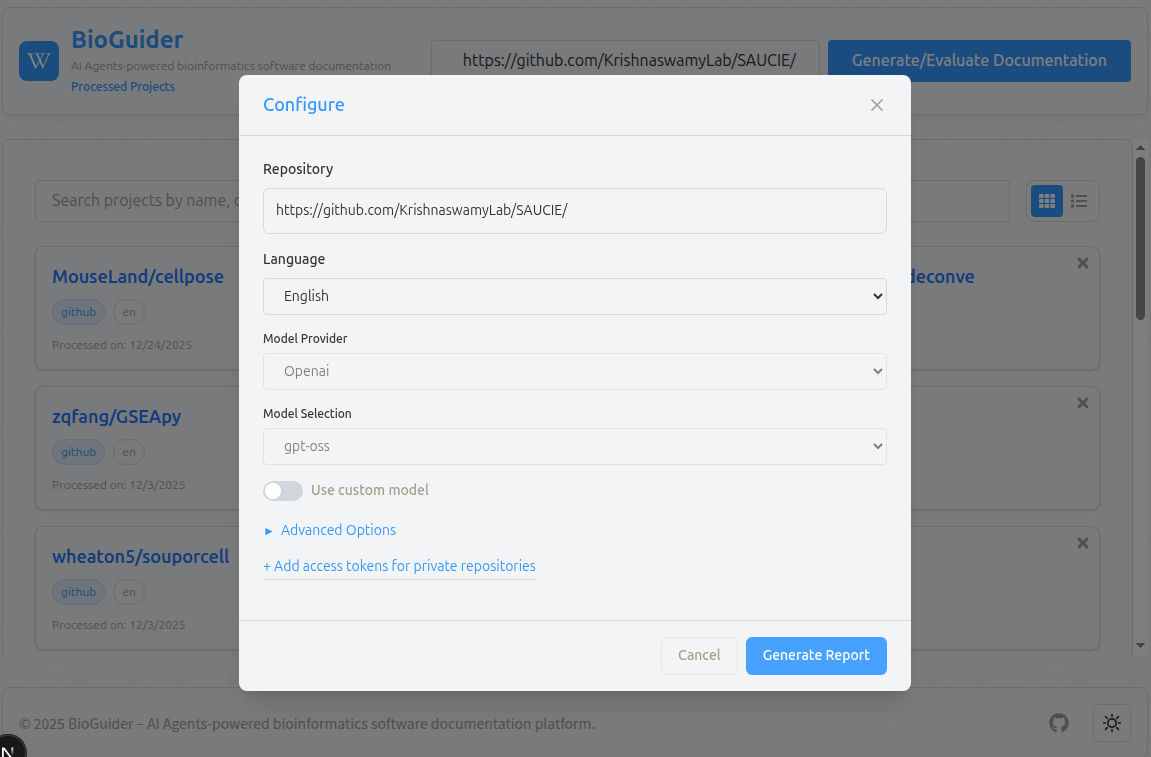

After clicking **“Generate Report”** in the Configure dialog, BioGuider initiates the automated evaluation workflow. During this stage, the system analyzes the repository structure and identifies relevant documentation components, such as the project overview, README files, Installation Section, User Guide/API, and Tutorial/Vignettes. A progress indicator is displayed to inform users of the current processing status and the documentation sections being evaluated in real time.

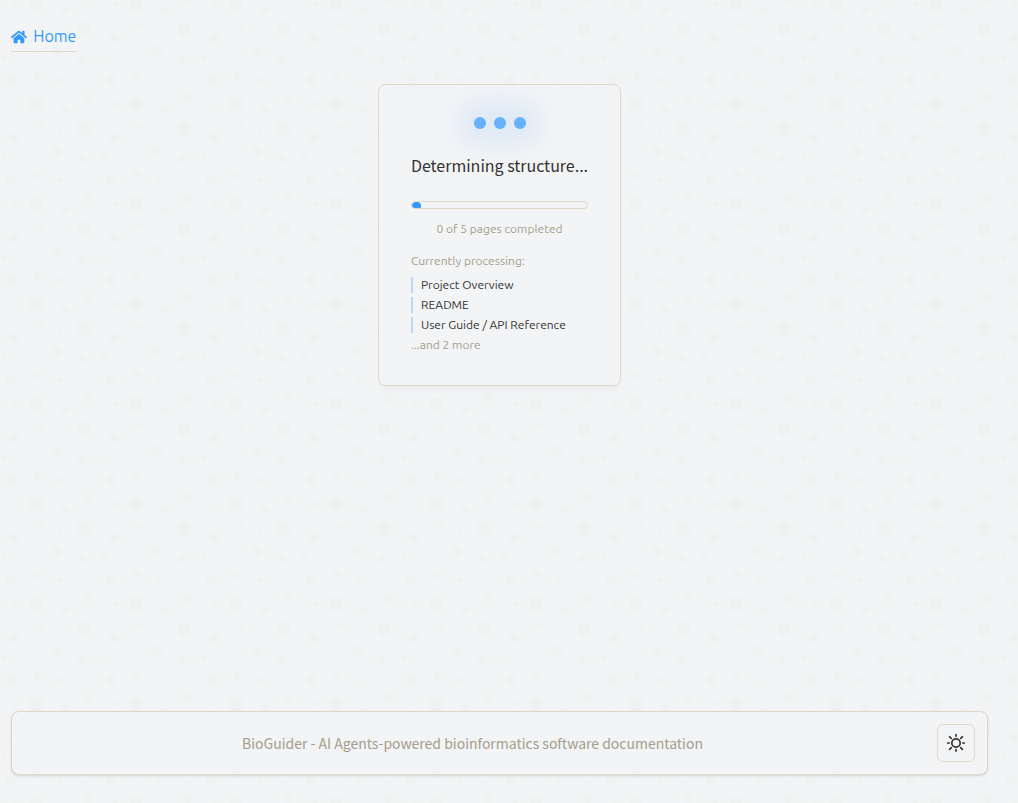

It will take a few minutes to generate the evaluation report. The report includes the following sections:

- Project Overview

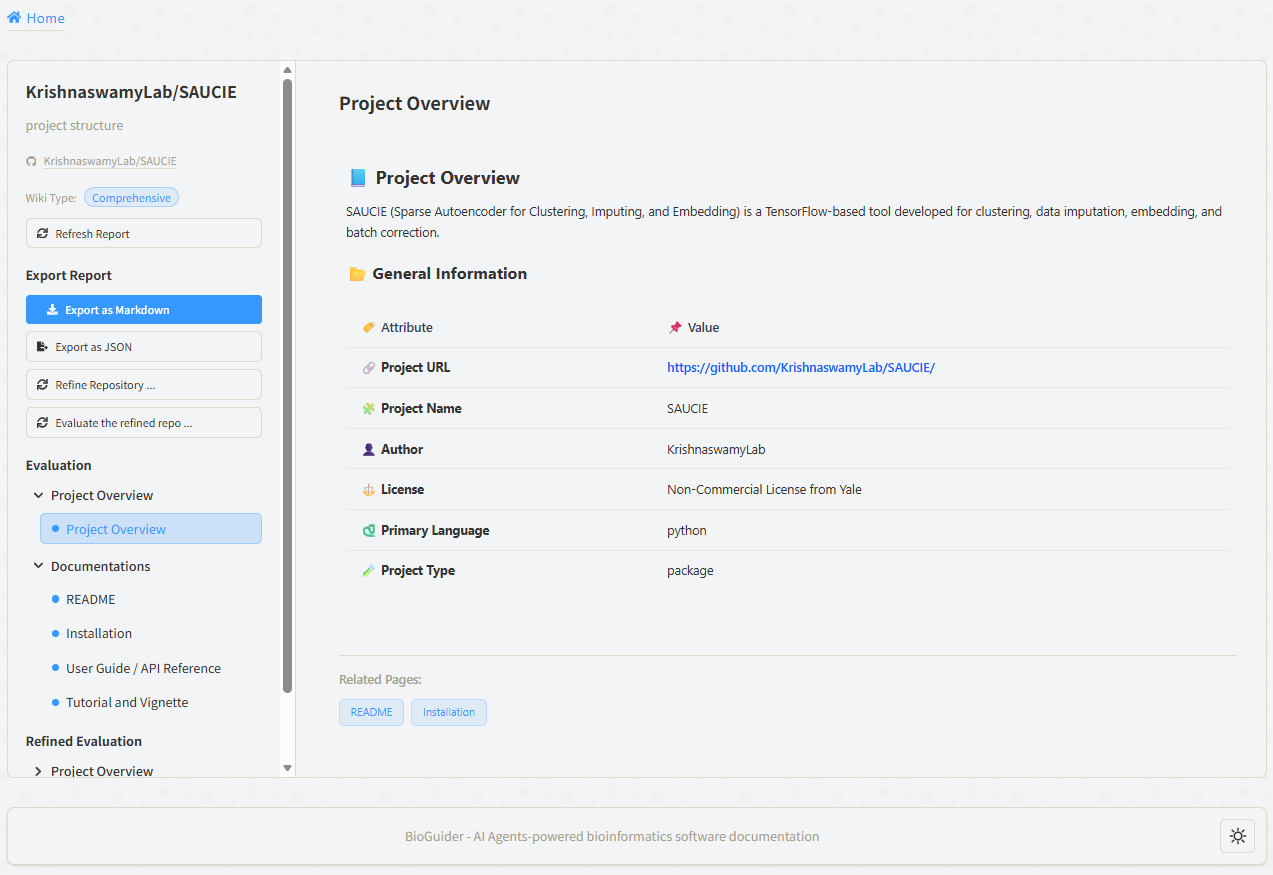

- Documentations

- README

- Installation

- User Guide / API Reference

- Tutorial and Vignette

The evaluation report can be downloaded from the left panel in Markdown (“Export as Markdown”) or JSON format (“Export as JSON”) for further analysis or archiving.

Each section under the **Documentations** will start with relevant scour files and their summary evaluation table. Even though the README file of SAUCIE received an overall score of 89, the readability (score 72) still has some room for improvement.

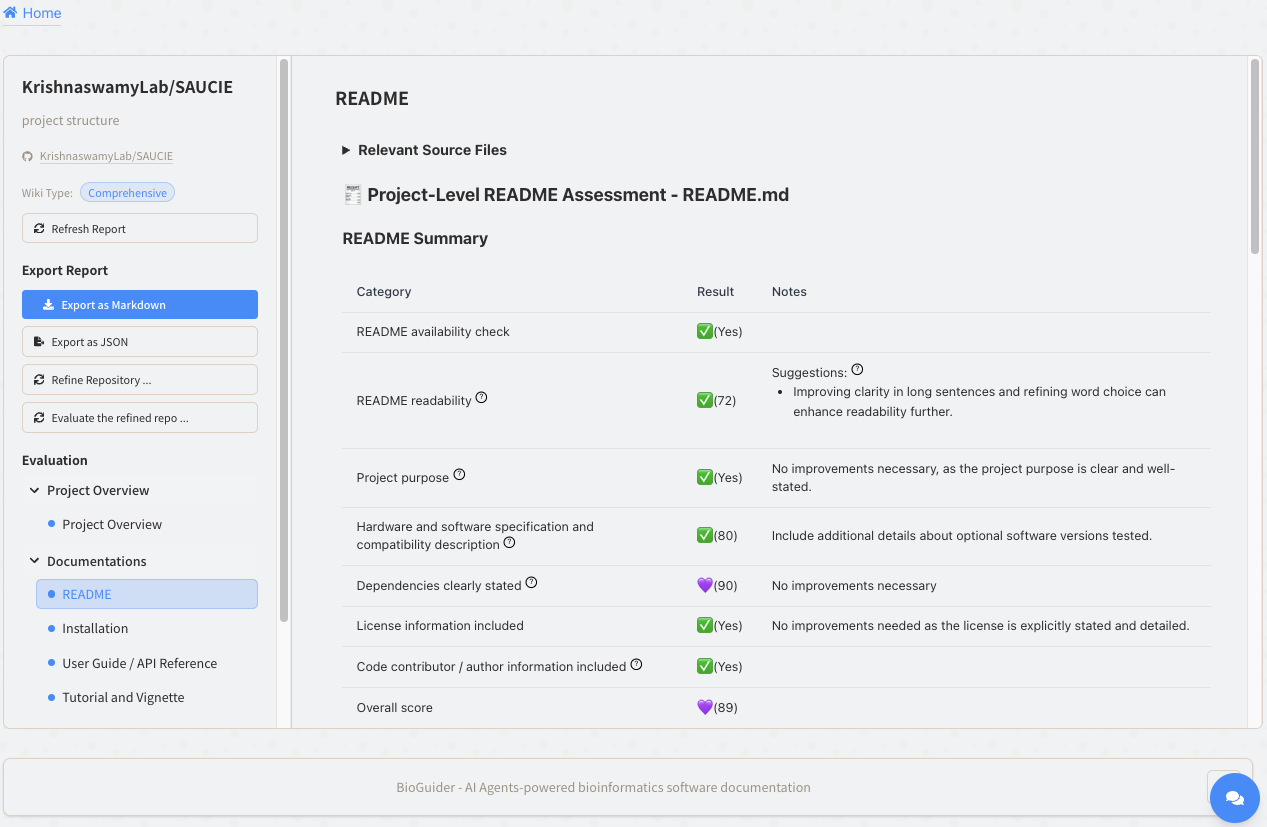

The detailed evaluation and suggestions can be found below the summary table. The BioGuider found three different types of readability errors in the README file and provided suggestions to increase the readability. In addition to readability, the BioGuider believed that the SAUCIE lacks deeper context about the project’s applications, importance, and target audience. Therefore, the BioGuider provides instructions for tool developers with formatted statements to improve the introduction of the tool. The BioGuider also evaluated the hardware and software compatibility and found that the README file does not discuss hardware specifications or compatibility with operating systems. This reminded developers to add more detailed hardware requirements to improve the success rate of installation.

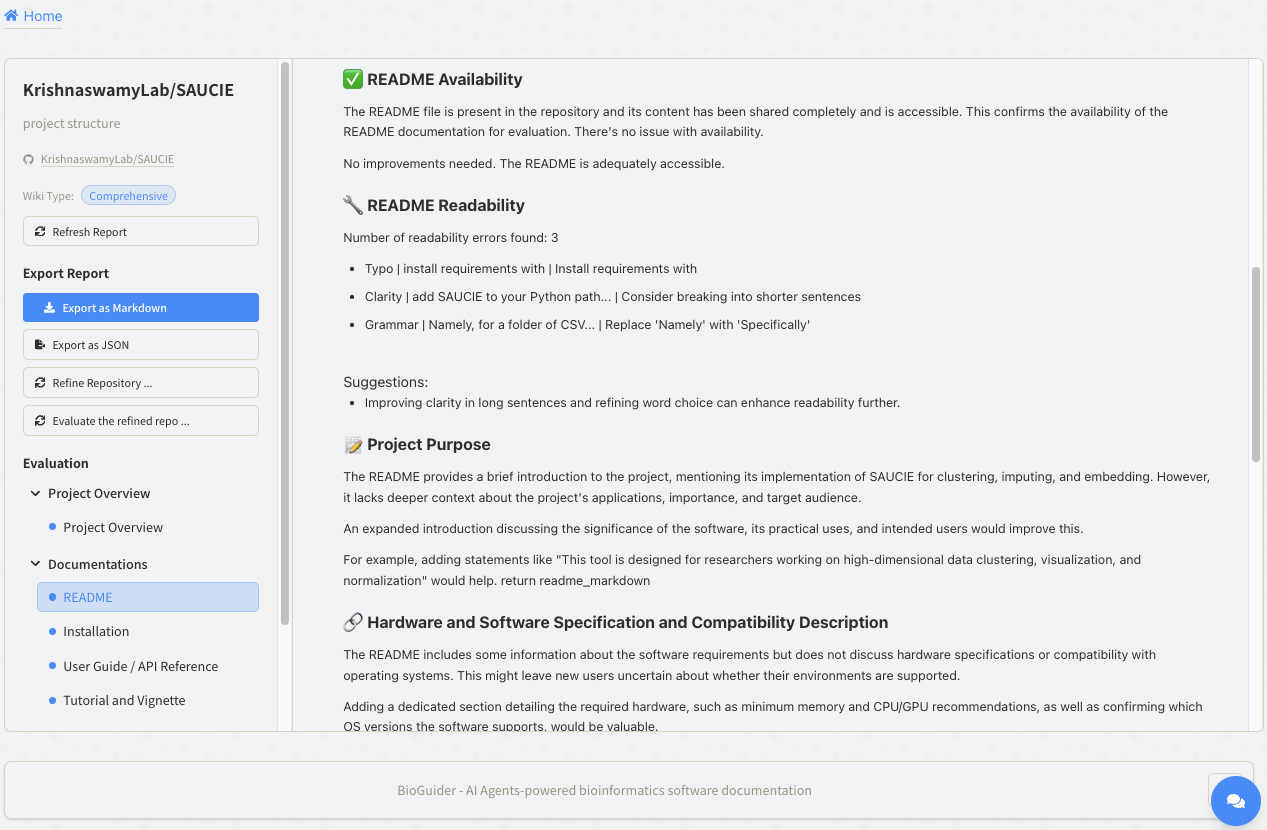

In the Installation page, the BioGuider provided an Overall score and many detailed sections to improve the success rate of installation, including the Step-by-Step installation Guide, Clarity of Dependencies, Compatible Operating System, and Hardware Requirements.

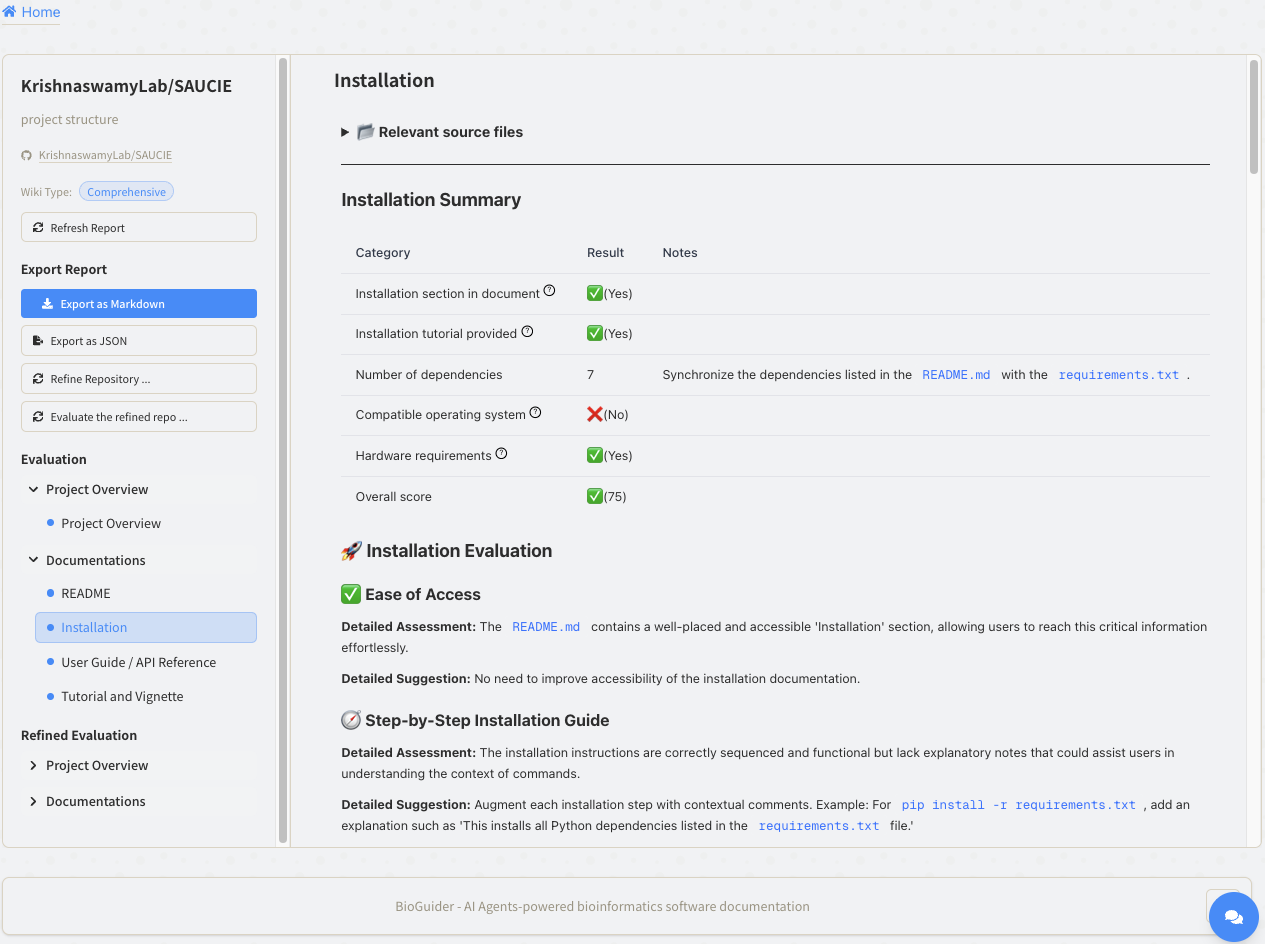

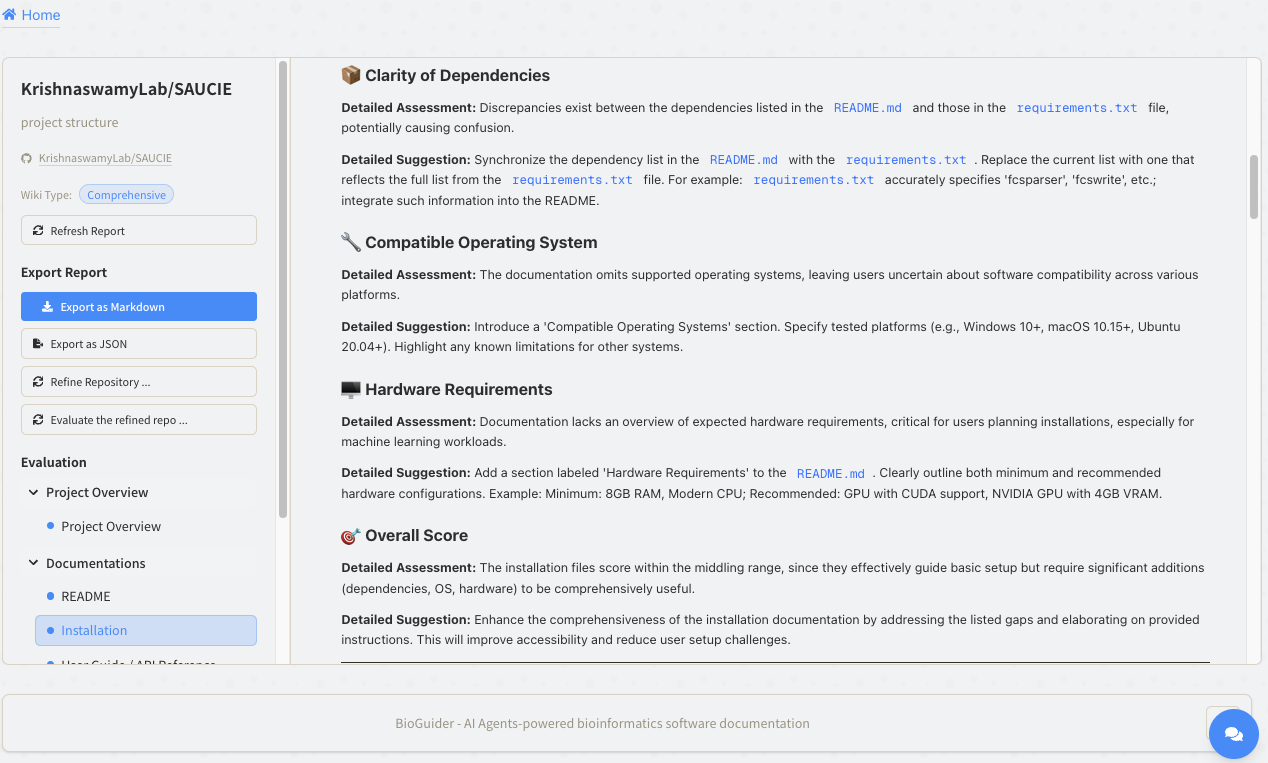

The BioGuider also evaluated the User Guide and Tutorial sections in the document. The reproducibility is one of the biggest challenges in the field of Bioinformatics. The higher quality of the User Guide / API reference and the Tutorial / Vignette documents can significantly increase the reproducibility. The BioGuider also evaluated consistency between documents and functional codes to ensure the reproducibility of the tool. The SAUCIE has a good quality of the User Guide with a score of 81, and the Tutorial with a score of 85. However, the BioGuider still provides some detailed suggestions after performing consistency evaluation. These suggestions help developers to review the concordance between documents and their functions’ parameters.

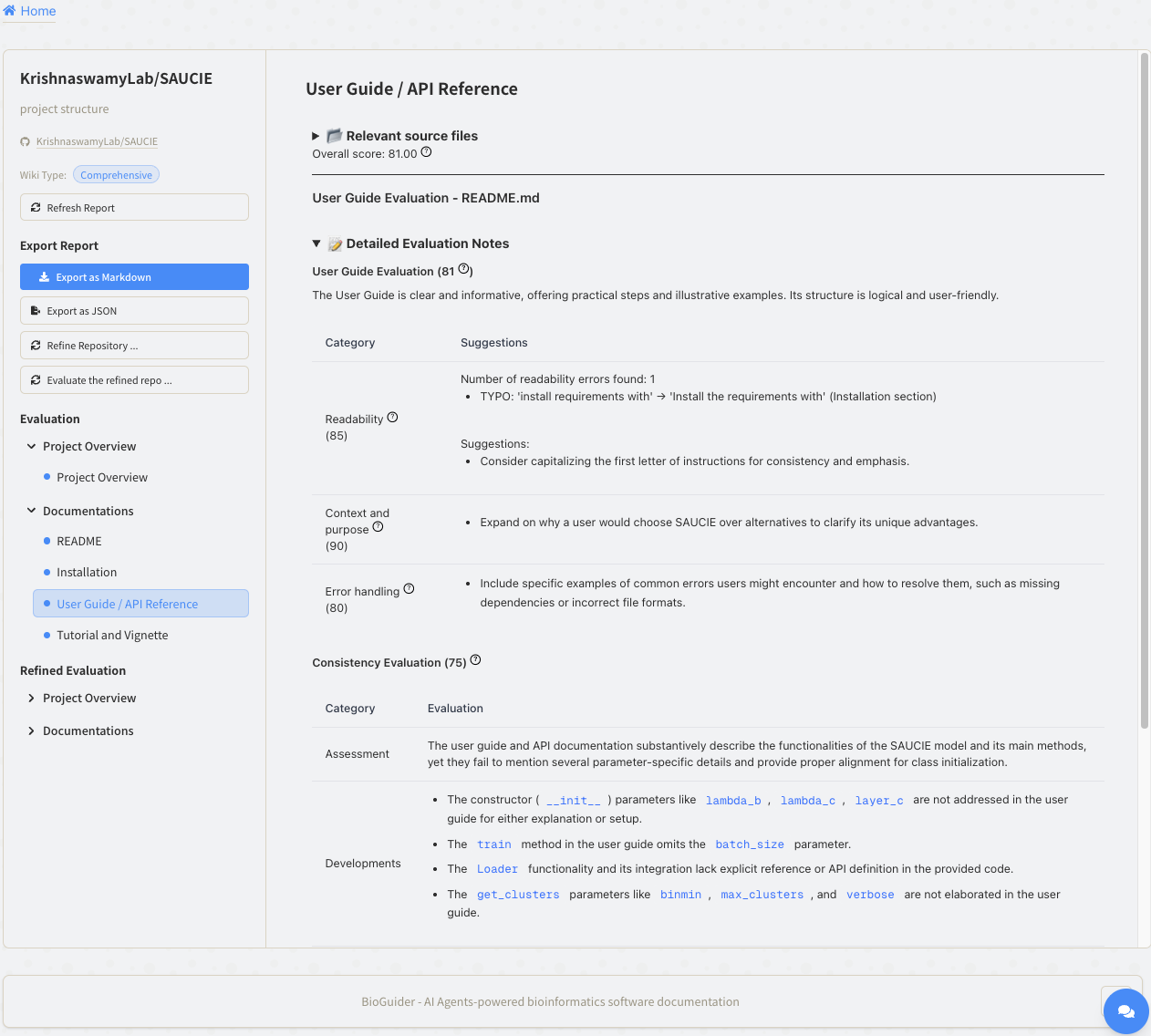

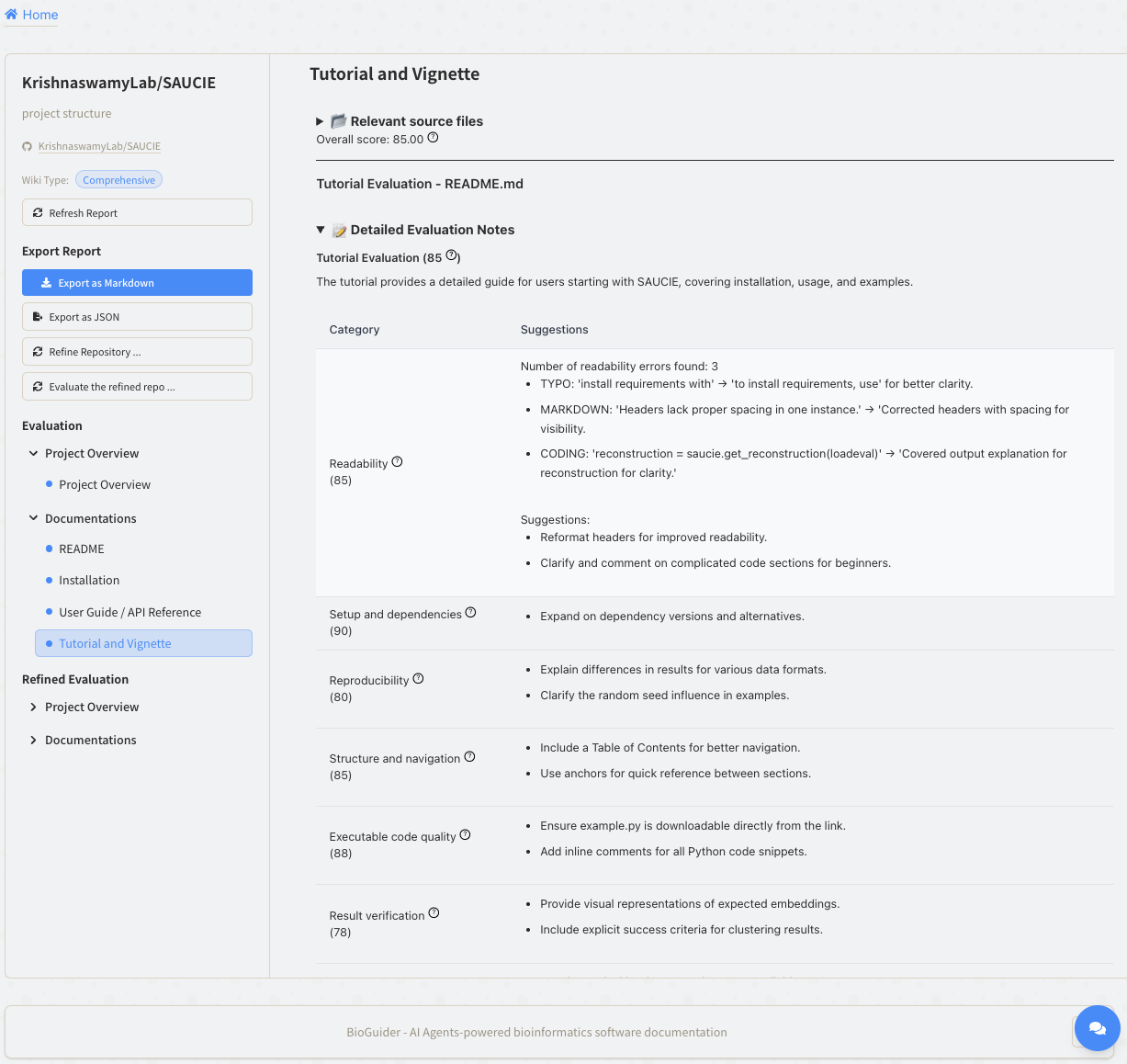

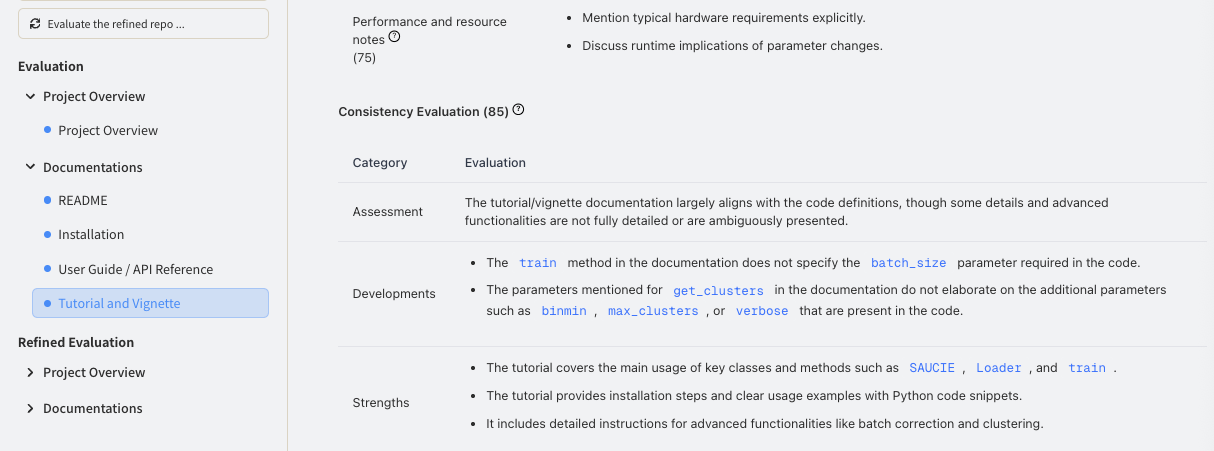

In addition to documentation evaluation, BioGuider supports automated documentation improvement. By clicking **“Refine Repository …”**, users can trigger a refinement workflow that generates improved versions of the identified documentation files. Once the refinement process is complete, the refined documents are automatically downloaded.

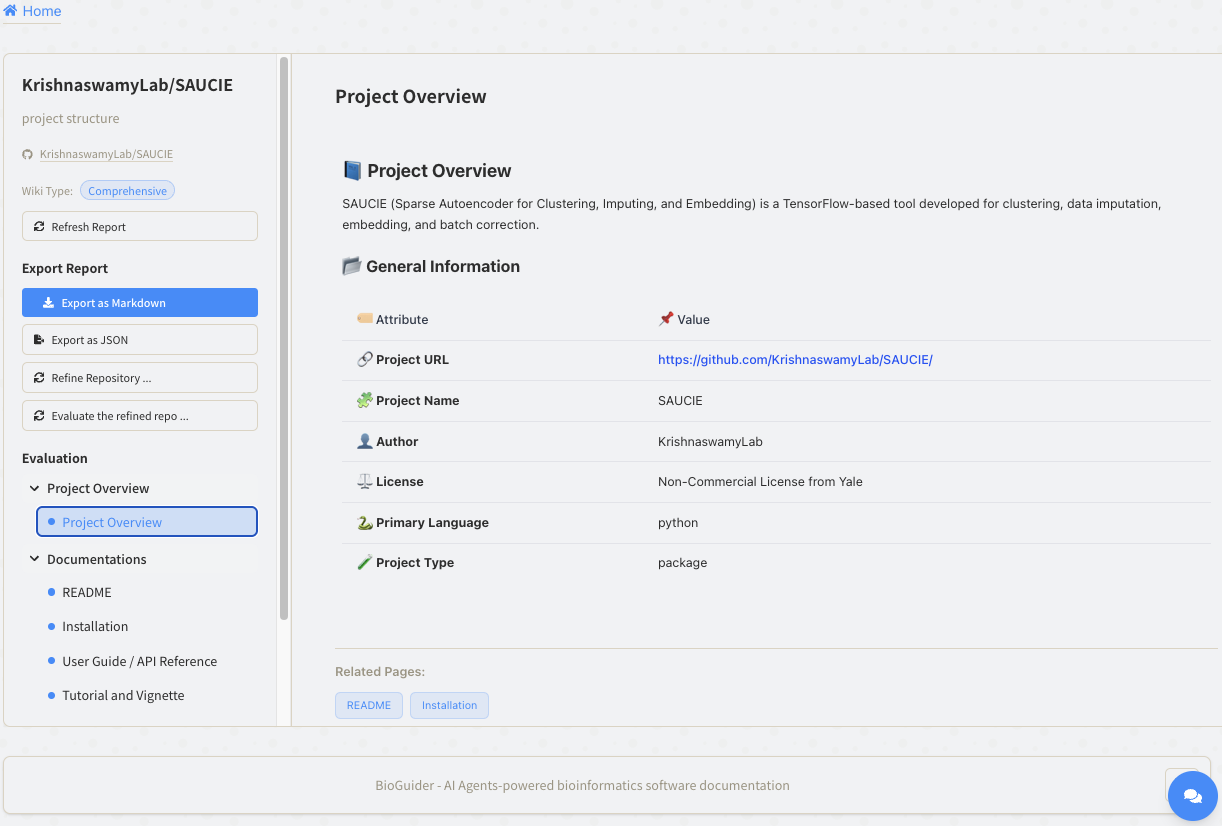

5 Click the Evaluate the refined repo button button

4 Click the Refine button

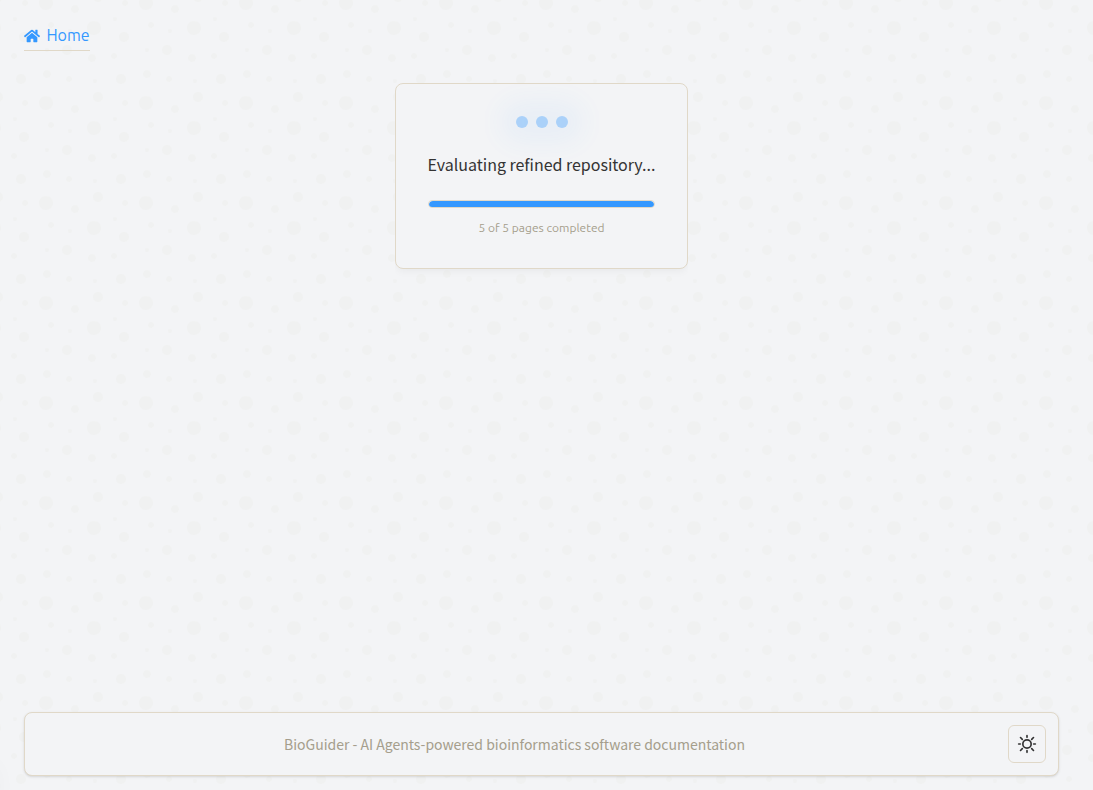

After refinement, users may optionally re-evaluate the improved documentation to assess changes in documentation quality (**“Evaluate the refined Repo …”**). The refined evaluation reports are stored in the project tree and can be accessed alongside the original results, enabling direct comparison between the initial and refined documentation assessments. We refined the original documents of SAUCIE and performed the second evaluation for the refined documents through the BioGuider. The BioGuider fixed the errors in the README file and improved the README readability from 72 to 95. The overall score of the README file was increased from 89 to 96.

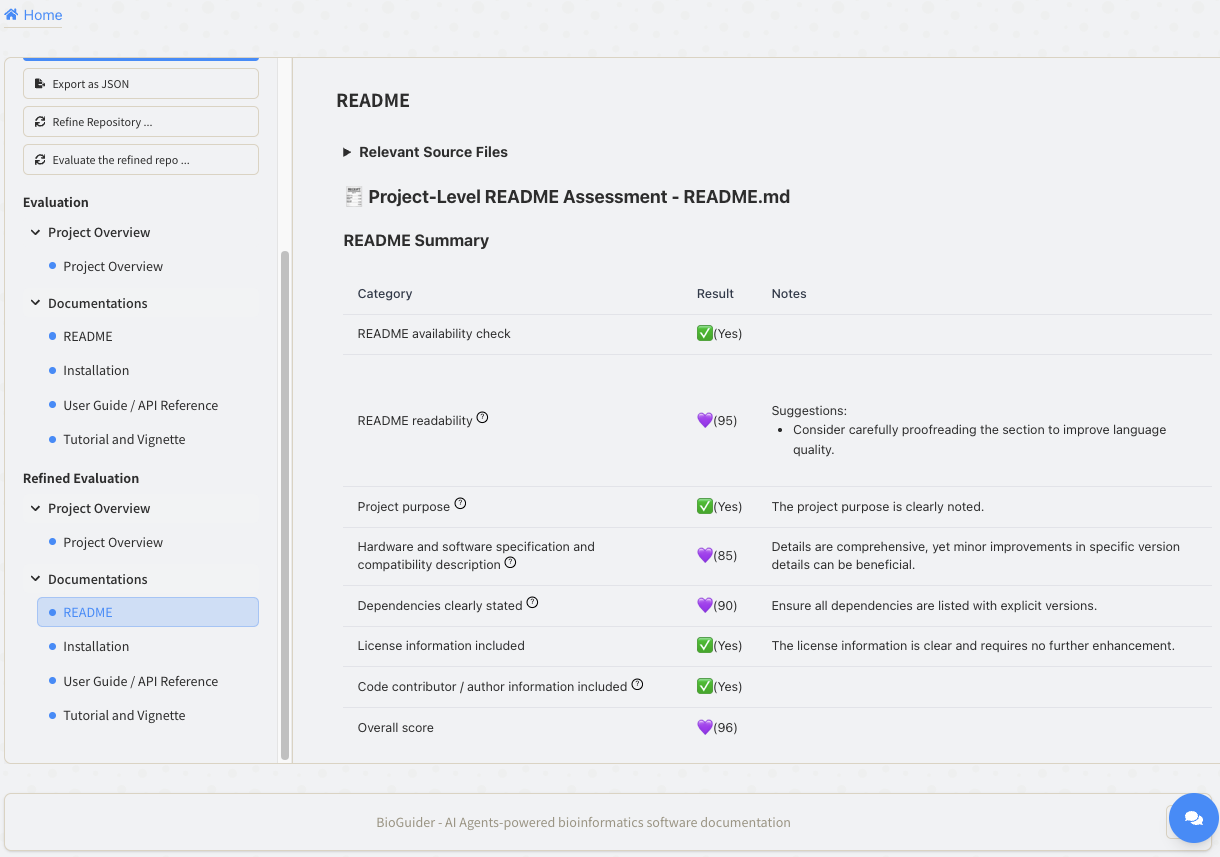

The second evaluation of the README file shows no error in readability. In addition, the project’s purpose, hardware specification and compatibility, and dependencies descriptions were improved.

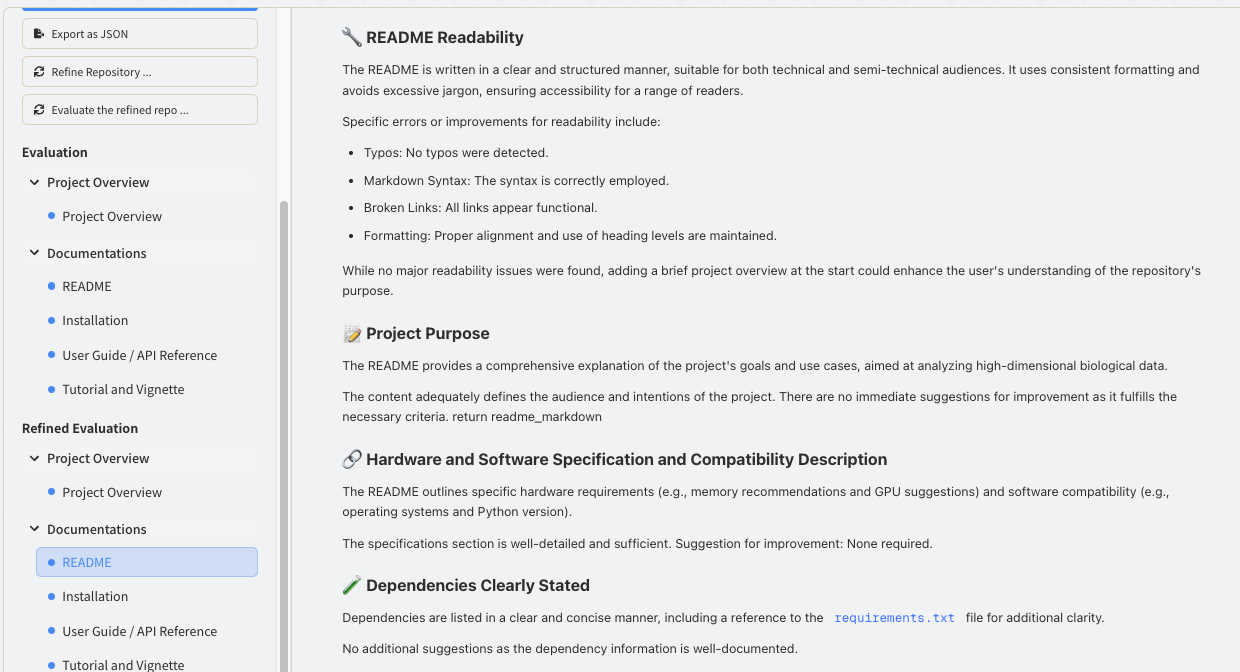

The Installation section’s overall score on the refined document was increased to 100. This indicates that BioGuider has successfully improved the installation guide.

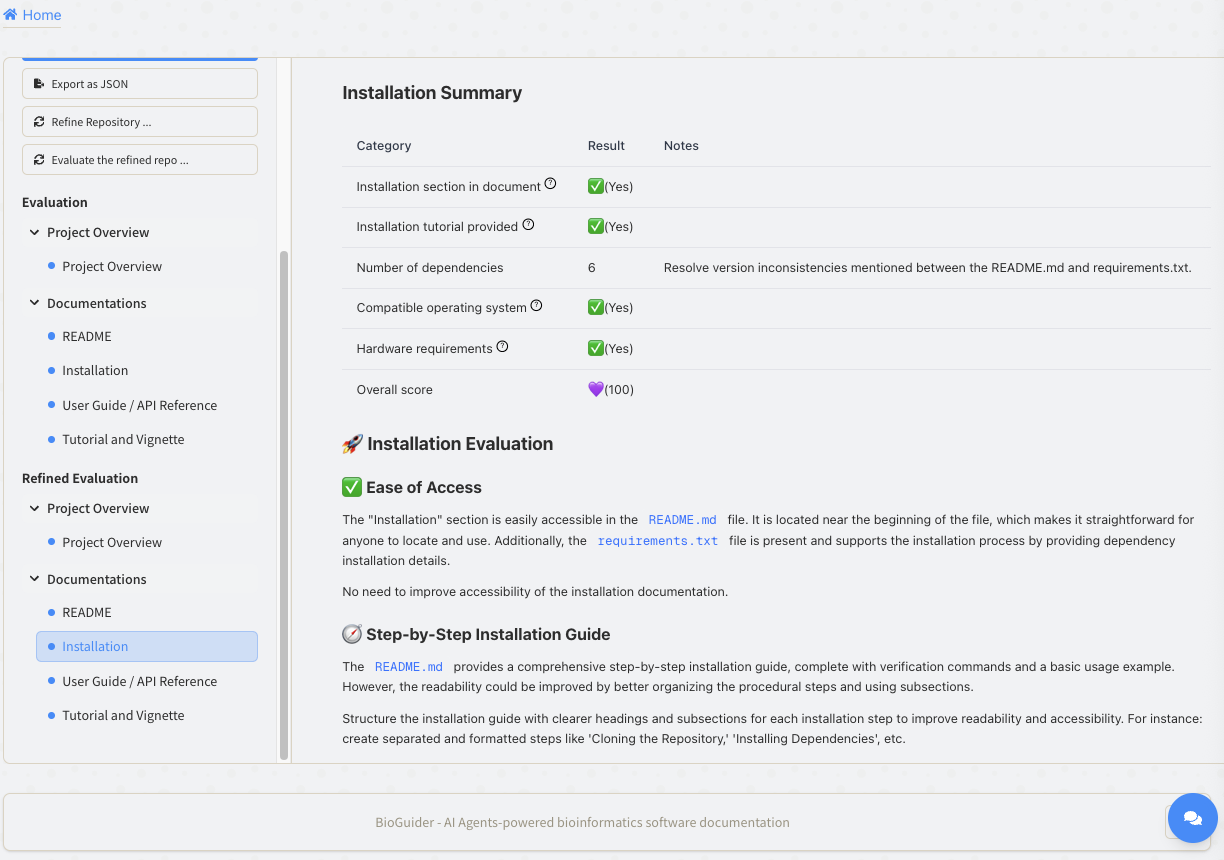

The evaluation score of the User Guide was increased from 81 to 89, especially on the Consistency Evaluation score (75 to 90). The suggestions from the functions’ parameters and documents were applied.

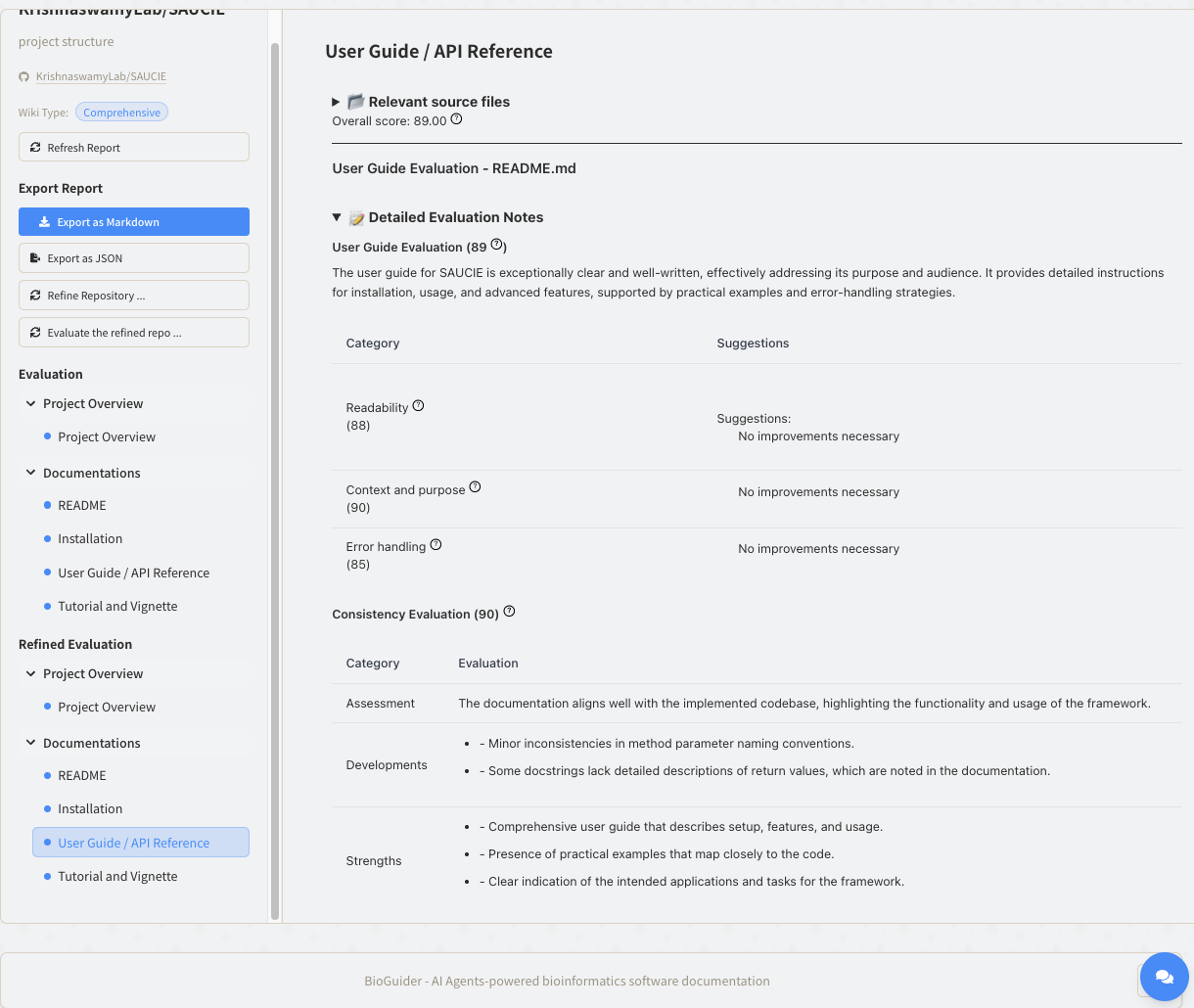

The tutorial section was slightly improved from 85 to 88. Even though the BioGuider applied some suggestions from the original evaluation, there are still some suggestions (for example, avoid absolute file path; give an example output visualization; include troubleshoot tip etc.) that were provided to the developers for improvement.

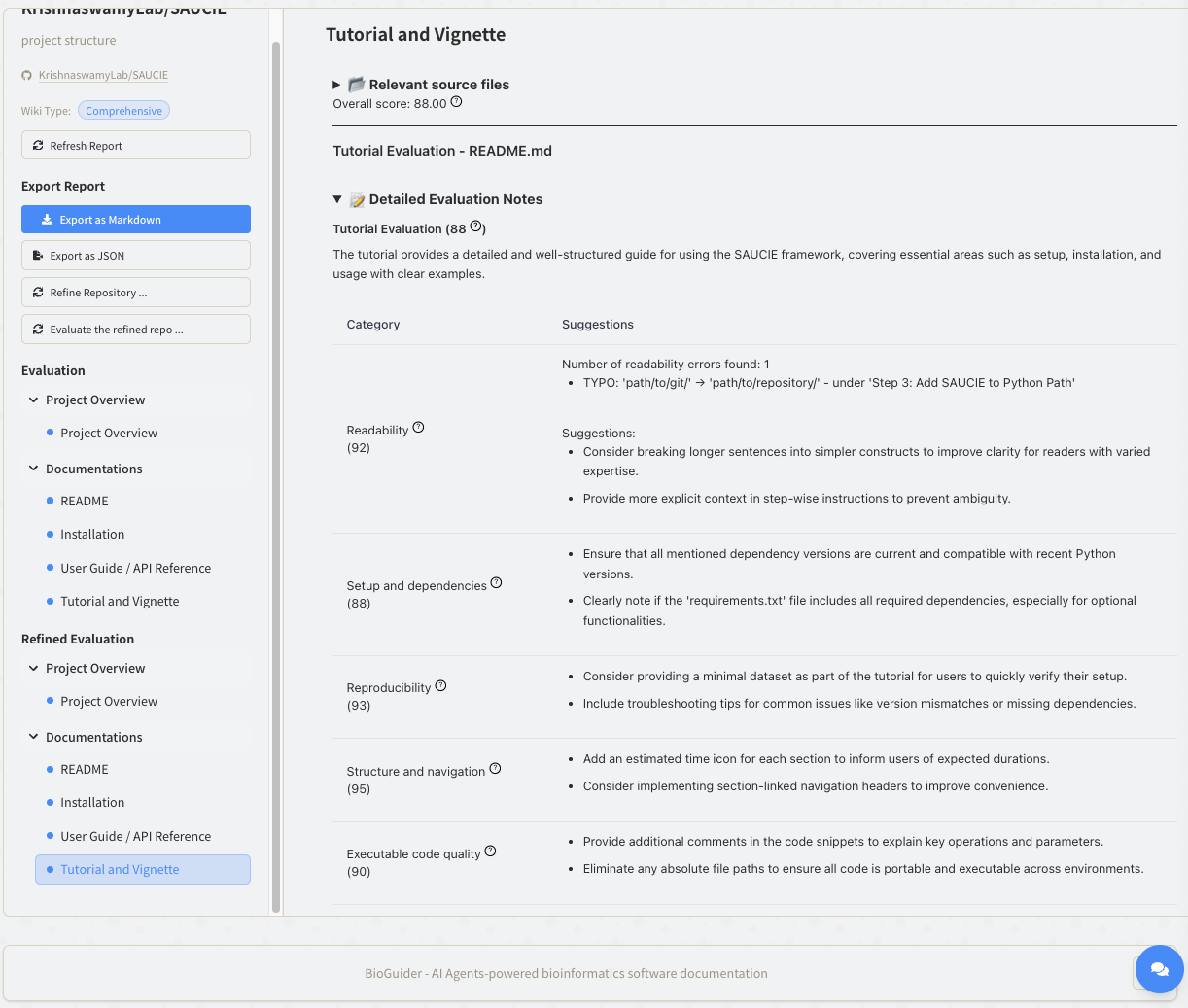

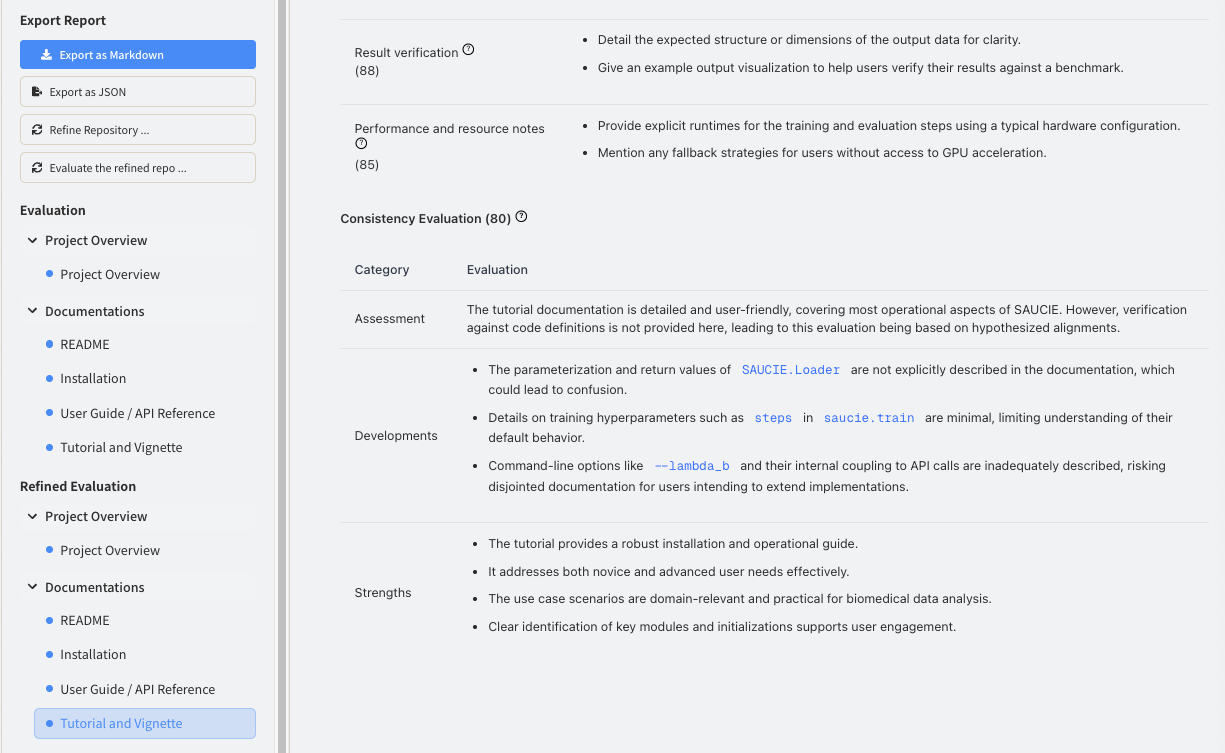

Overall, the BioGuider’s multi-AI agents evaluated and refined the original documents of SAUCIE. Through the scores and detailed suggestions that the BioGuider provided, the developers could create high-quality documentation for the tool. At the end, the original and refined documents of SAUCIE were provided below.

**Example comparison of SAUCIE document before and after BioGuider correction**

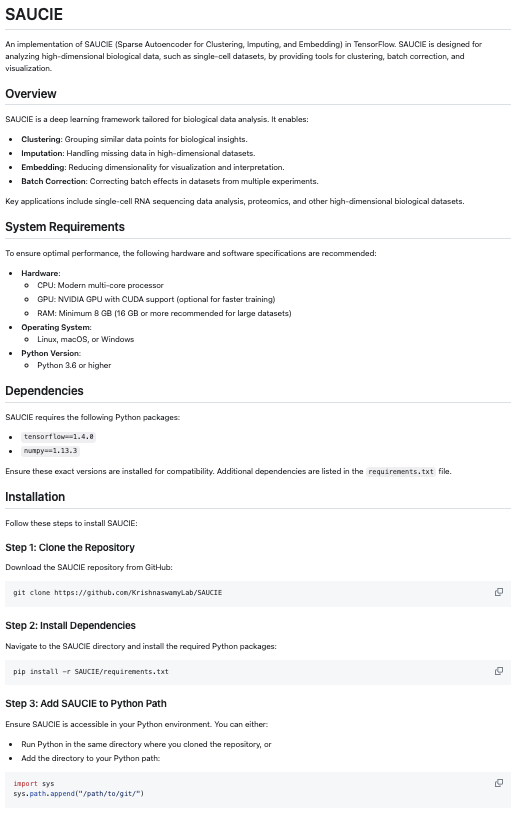

The original SAUCIE README file.

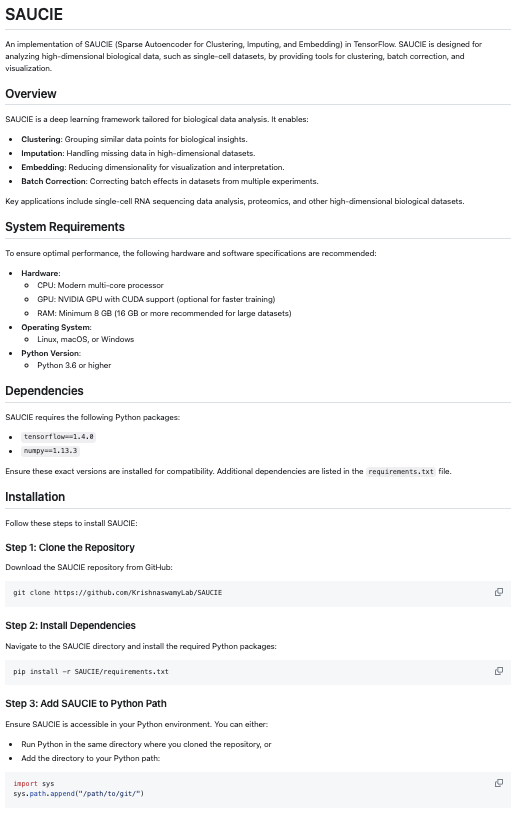

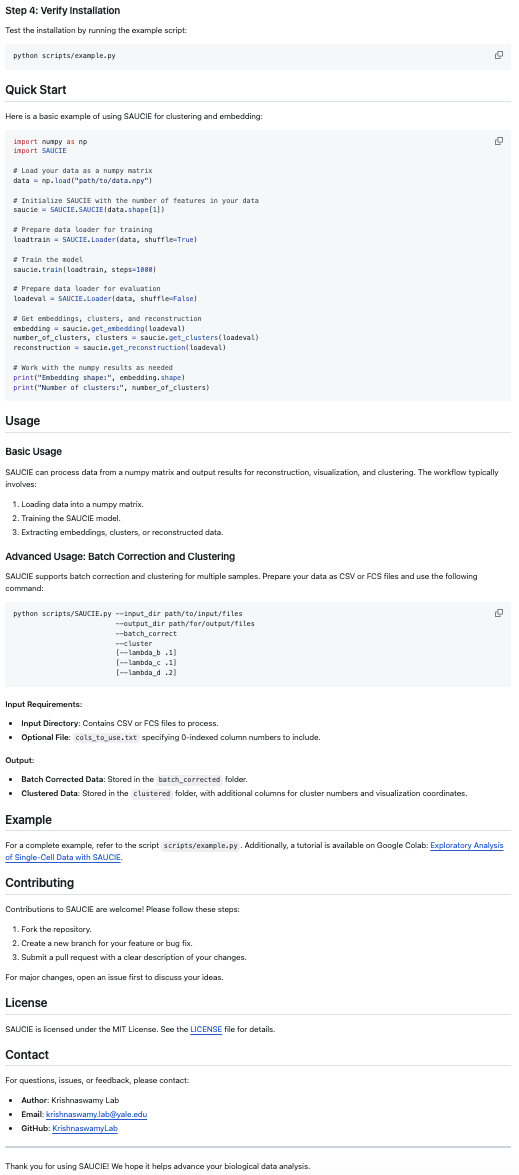

The revised SAUCIE README file.
